# Development of a differential panel to identify *Peronospora belbahrii* races in sweet basil (*Ocimum basilicum*)

**DOI:** 10.1101/2025.09.18.674471

**Authors:** Kelly S. Allen, Alexander J. Barrett, Yariv Ben-Naim, Robert Mattera, Christian A. Wyenandt, Yigal Cohen, James E. Simon, Li-Jun Ma

## Abstract

First reported in the United States in 2007, downy mildew of sweet basil (*Ocimum basilicum*), caused by the oomycete pathogen *Peronospora belbahrii*, has become a major challenge in global basil production. Downy mildew-resistant cultivars have been developed and commercially available since 2018, but observations of isolates overcoming resistance were reported after each new introduction. The rapid development of pathogens overcoming genetic resistance suggests the evolution of new races under selective pressure. In this study, we report on the development of a differential panel of distinct basil cultivars with varying and differential sources of genetic resistance tested against *P. belbahrii* strains obtained from geographically diverse multi-year collections from basil-producing locations in the US, Israel, and Italy. Screening multiple *P. belbahrii* isolates revealed three distinct virulence profiles that distinguished race 0, race 1, and race 2. Races 1 and 2 were defined by virulence in cultivars with single gene resistance loci *Pb1* and *Pb2*, respectively. Evaluation of cultivars with quantitative resistance revealed more variation in susceptibility or partial resistance, suggesting “broad spectrum” (or race-nonspecific) resistance. With continued isolate collection and screening efforts, the denomination of *P. belbahrii* races may be further expanded, and newly developed resistant cultivars can be added to the proposed panel to definitively evaluate novel *P. belbahrii* strains. This study lays the groundwork to further explore dynamics of race evolution and mechanisms of resistance breakdown, which will be essential for discovering new sources of resistance for controlling basil downy mildew.

## Introduction

Basil downy mildew is one of several downy mildew diseases impacting culinary herb production globally, which can rapidly result in total crop loss in susceptible varieties with estimated economic impacts in the tens of millions of dollars (Hoffmeister et al. 2020; Roberts et al. 2009; Wyenandt et al. 2015). Basil downy mildew (BDM) is caused by *Peronospora belbahrii*, a biotrophic oomycete pathogen that infects basil leaves by directly penetrating the epidermal leaf tissue or entering through leaf stomatal openings (Cohen et al. 2017). Symptoms and signs of infection include: interveinal chlorosis and gray, mildew-like sporulation on the underside of leaves (sometimes emerging on the upper surface as well) (Belbahri et al. 2005; Garibaldi et al. 2007; Koroch et al. 2013). Sporangiophores, the spore-bearing structures that emerge from stomata of infected leaves, produce abundant sporangia which are wind-dispersed in both field and greenhouse production settings (Cohen et al. 2017).

In the US, the BDM population is known to survive in southern FL and northern Mexico, and spreads northward due to natural weather patterns and/or distribution of pre-symptomatic plants through nursery stock (Wyenandt et al. 2015). As basil crops are planted in the field during late spring and summer months up the East Coast, typical patterns of aerial dispersion have been observed as primarily late-season infections following earlier activity in southern regions (Wyenandt et al. 2015). The US domestic and global populations of BDM have not yet been thoroughly evaluated in genetic studies, so it remains unknown how genetically diverse active field isolates (populations) may be across years and locations.

Sweet basil (*Ocimum basilicum*) is an outcrossing allotetraploid with limited genomic resources, and breeding new fertile cultivars that are disease resistant and retain desired horticultural traits is challenging (Ben-Naim et al. 2015, 2018; Pyne et al. 2014, 2015). Bar-Ilan University in concert with “Genesis Seed LTD” developed the Prospera resistant lines family with qualitative (major gene-mediated) resistance by introgression of highly resistant *O. americanum* accessions with susceptible Sweet Basil (Genesis Seed) (Ben-Naim et al. 2018).

Resistance in the Prospera lines is linked to a pair of dominant loci *Pb1A* and *Pb1A’* (Ben-Naim et al. 2018; Ben-Naim and Weitman 2022). In 2024, the Prospera Active series was released with a new source of genetic resistance derived from a gene/locus referred to as *Pb2* (Ben-Naim and Weitman 2022). These lines are resistant to race 0 and race 1 downy mildew isolates with or without the combination of *Pb1* (Ben-Naim and Weitman 2022). Other resistant cultivars released include Amazel Basil (Proven Winners, USA), which is a sterile hybrid with qualitative resistance derived from the same *O. americanum* accession as the Prospera lines, and the recent introduction Pesto Besto (Proven Winners, USA), which carries the same source of resistance as Amazel Basil but can be grown from seed (Ben-Naim and Weitman 2022; McGrath 2024). In addition, Enza Zaden cultivar Eleonora offers intermediate resistance and has exhibited partial control in several trials (McGrath 2024).

Sweet basil Mrihani [*O. basilicum* (Horizon Seed Co., Williams, OR)] was identified as a BDM-resistant cultivar in the basil breeding program at Rutgers University through a screen of 39 basil accessions representing a broad geographic distribution (Pyne et al. 2014, 2015). A quantitative trait locus (QTL) analysis was used to identify one major QTL (dm11.1) and two minor QTL (dm9.1 and dm14.1) conferring downy mildew resistance derived from a cross with Mrihani (Pyne et al. 2017, 2015, 2018; Simon et al. 2018). Mrihani was first suspected to be a separate *Ocimum* species, but genetic analysis showed it was an *O. basilicum* and thus it was incorporated into the Rutgers breeding program as a DMR (Downy Mildew Resistant) parent. The Rutgers breeding program produced four DMR sweet basil cultivars from Mrihani crossed with BDM-susceptible Newton (SB22) (Simon et al. 2018). These cultivars, Rutgers Devotion DMR, Rutgers Obsession DMR, Rutgers Passion DMR, and Rutgers Thunderstruck DMR, have offered partial control of downy mildew with apparent broad-spectrum, non-race-specific resistance in prior field studies (McGrath 2024). Linkage mapping performed in the Mrihani x SB22 F_2_ mapping population suggested the action of one or more dominant genes controlling BDM resistance (Pyne et al. 2017).

The introduction of resistant cultivars has predictably been accompanied by various reports of a “breakdown of resistance”, or more accurately, novel pathogen strains overcoming resistance across geographic locations (Ben-Naim and Weitman 2022; Browder 1980; Feng et al. 2018a; Habgood 1970; van der Plank 1969). The terms “strain”, “pathovar”/“pathotype”, and “race” may sometimes be used interchangeably, but all typically refer to the subspecific classification of pathogens based on unique colonization of host genotypes (Schulze-Lefert and Panstruga 2011). The concept of physiological races is generally understood as an isolate or group of isolates that exhibit the same virulence pattern across a set of host lines within a species (Browder 1980; Edel-Hermann and Lecomte 2019; Schulze-Lefert and Panstruga 2011). Pathogen races should therefore be demarcated by evaluating multiple isolates representing diverse locations, hosts, seasons and years, establishment of a standardized panel of differential cultivars or lines, and replicated controlled testing of all isolates against the panel (Browder 1980; Edel-Hermann and Lecomte 2019; Schulze-Lefert and Panstruga 2011).

The development of new phytopathogen races overcoming host resistance has been documented in lettuce downy mildew, spinach downy mildew, and other economically important pathosystems in response to the introduction of resistant cultivars (Feng et al. 2014, 2018b, 2018a; Irish et al. 2003; Lebeda and Reinink 1994; Parra et al. 2016). The first spinach downy mildew (*Peronospora effusa*) race overcoming a resistant cultivar was reported in the US and Europe in 1824 (Irish et al. 2003). In 1996, there were five races identified and by 2018, nineteen distinct races with differential virulence signatures on spinach cultivars have been denominated by the International Working Group on *Peronospora* in spinach (IWGP) (Feng et al. 2018b; Irish et al. 2003). Lettuce downy mildew, caused by *Bremia lactuceae*, has similarly developed multiple races overcoming resistant cultivars (Parra et al. 2016, 2021). The International Bremia Evaluation Board (IBEB) has reported forty-one discrete races of *B. lactucae* historically in the US and Europe, though contemporary races are fewer as they are no longer active (International Bremia Evaluation Board (IBEB) n.d.). Although BDM is mainly a concern for basil, other members of the *Lamiaceae* family besides *O. basilicum* serve as hosts for *P. belbahrii*, thus possibly aiding its spread and survival (Ben Naim et al. 2019; Cohen et al., 2017). *P. belbahrii* was observed in several *Salvia* spp., *Rosmarinus officinalis*, *Nepeta curviflora*, *Micromeria fruticosa*, and *Agastache* spp. in Israel (Ben Naim et al. 2019). The colonization of these Lamiaceae species may play a role in the epidemiology, prevalence or even the development of new strains (Ben Naim et al. 2019). More work is required to understand the host range of *P. belbahrii* across different geographic regions in agricultural and wild hosts.

Until 2021, *P. belbahrii* race evolution was anticipated but not reported. In 2021, two races of *P. belbahrii* were reported in Israel based on the differential virulence of the isolates infecting basil with two different sources of resistance (Ben-Naim and Weitman 2022). This initial description of *P. belbahrii* races was reported after screening two isolates against a limited number of differential lines, and simple-sequence repeat (SSR) profiling of multiple isolates supported the separation of isolate groups based on unique molecular signatures (Ben-Naim and Weitman 2022). Apart from this study, cultivar trials and germplasm screenings are often performed in open field settings using natural inoculum, with disease incidence and severity assessed weekly (Ben Naim et al. 2025; McGrath 2024; Patel et al. 2021). In many cases, these trials do not include a full panel of all available resistant cultivars, use differential scoring criteria, confounding the identification and detection of new races of *P. belbahrii*.

Additionally, natural inoculum can be highly variable from season to season. In order to conduct precise single-isolate screens, environmental and experimental controls are required, and necessitate the need for standardization among research groups.

The objective of this research was to establish and report a standardized differential panel for the identification and separation of novel *P. belbahrii* isolates into discrete races using a reproducible controlled inoculation method. The proposed differential set includes several resistant lines that have the potential to differentiate existing *P. belbahrii* races, and includes new resistant lines that may be useful in helping to identify new *P. belbahrii* races. As a major limitation of working with downy mildews and other obligate pathogens is the proper handling and sequestering of multiple isolates, we additionally describe best practices for maintaining *P. belbahrii* isolates in long-term storage for the retention of viable spores.

## Materials and Methods

### Basil propagation

Basil cultivar seeds were sown into soilless growing media (Promix BX Premier Horticulture Ltd., Qc) in standard plug flats and thinned to one seedling per plug after 1-2 weeks. The full panel of plants were transplanted to two 24-cell trays with four to six plants per cultivar for inoculations. Plants were maintained under controlled-environment greenhouse conditions at the University of Massachusetts CNS Research and Education Greenhouses located in Amherst, MA (24°C (75°F), 14/10 hour Day/Night cycle, supplemental lighting of half metal halide and half high-pressure sodium lights) and the Rutgers New Jersey Agricultural Experiment Station Research Greenhouse located in New Brunswick, NJ (23-27⁰C, 16/8hr Day/Night cycle, ∼100µmol/m²/s supplemental lighting). Plants were regularly hand-watered and fertilized 3 times weekly during irrigation with Peters Professional 20-10-20 peat lite fertilizer at 200 ppm nitrogen (UMass) or weekly with a standard 20-20-20 fertilizer mix (Rutgers).

### *Peronospora belbahrii* isolate collection and maintenance

Downy mildew isolates (n = 9) were collected from research plots, private garden centers, production fields and greenhouses from 2020-2024 in the US, Israel, and Italy. Symptomatic whole plants or cut stems were taken from each location, with a single location representing one isolate.

Sporulation was forced by incubating symptomatic plant material in a high humidity chamber (RH ≥96%) on a greenhouse bench or in a growth chamber for 8-12 hours with 10 hours of darkness to ensure sporulation (Cohen et al. 2017). Freshly sporulating leaves were collected and placed in chilled (4°C) sterile water, vortexed, and then filtered through a double layer of cheesecloth to remove debris. The sporangial suspension was then re-filtered through quadruple-layered cheesecloth to further remove sporangiophores, soil particles, and other debris. The suspension was quantified using a hemocytometer. Sporangia suspensions of 1-5×10^4^ sporangia/mL were utilized for isolate maintenance by spray inoculating plants until run-off and incubating in a humidity chamber kept in dark conditions for 12-24 hours. Isolates were maintained on the genotype from which they were originally isolated. Healthy plants were inoculated every week to build sufficient inoculum for differential screening. Only one isolate was actively maintained per greenhouse or growth chamber, at a time. If other isolates were collected, they should be contained in different greenhouse, and sporulation and inoculations were staggered to reduce the possibility of cross-contamination.

### Storage and recovery of viable *P. belbahrii* inoculum

Once isolates were re-inoculated successfully onto their host genotypes, spores were collected during maintenance inoculations for long-term storage of viable spores. Freshly incubated leaves with heavy sporulation were collected and lightly pressed against sterile P5 filter paper, rolled and placed into labeled 50mL conical tubes and frozen at −20°C (and moved to −80°C for longer-term storage), or stored directly at −80°C. Regeneration of frozen spores was performed by removing and unrolling frozen leaves after 2 hours of defrosting at 4°C or 20 minutes at RT, and removing sporangia from the leaf tissue by gently spraying with sterile water into a vessel on ice. Recovered sporangial suspension was then filtered and inoculated onto intact plants as previously described. Several conditions for long-term storage and recovery of viable material were tested, and viability was confirmed by quantifying *in vitro* germination as a percentage of 300 randomly sampled sporangia on 1.5% water agar incubated for 24 hours at 10-12°C, as well as *in vivo* plant inoculation (Supplementary Table 1).

### Basil differential panel selection and screening

Basil cultivars were selected for differential panel screening based on prior performance in greenhouse and field trials and representing commercially available cultivars in use in production systems. Cultivar Superbo is an Italian Genovese-type variety developed by SAIS Sementi (Cesena, Italy) that is susceptible to downy mildew. Rutgers DMR cultivars Devotion, Passion, and Obsession were selected as commercially-available, QDR accessions. Resistant cultivars with qualitative (vertical resistance) included Prospera PS5 or Prospera CG2 (*Pb1*), Prospera Active Noga (*Pb2*), and Prospera Active Mia (*Pb1/Pb2*). Additionally, two lines of basil representing distinct species were included, US86 (*Ocimum tenuiflorum*, PI652056) from the USDA Basil Germplasm Collection, and KL3 (*Ocimum americanum*), a Rutgers-coded germplasm accession.

Four to six plants per genotype tested were included per block in a randomized complete block design across two to three humidity chambers (Supplementary Figure 1). Plants were grown to first or second true leaf set expansion, approximately three to four weeks old.

### Evaluation and rating basil downy mildew disease symptoms

Disease incidence (DI) and disease severity (DS) scores were taken from each plant. DI was rated as a percentage of true leaves with visible sporulation on affected plants (McGrath 2020). The first one to two leaf pairs were examined for sporulation by eye and/or under a magnifying lamp (any axillary leaves present were not included as they may not have emerged at the time of inoculation). The DS score was assigned using an ordered categorical scale in which 0=no sporulation, 1=low sporulation density (1-10% leaf area coverage), 2=moderate sporulation density (11-40%), 3=high sporulation density (41-100%) (Figure 1).

**Figure 1.**
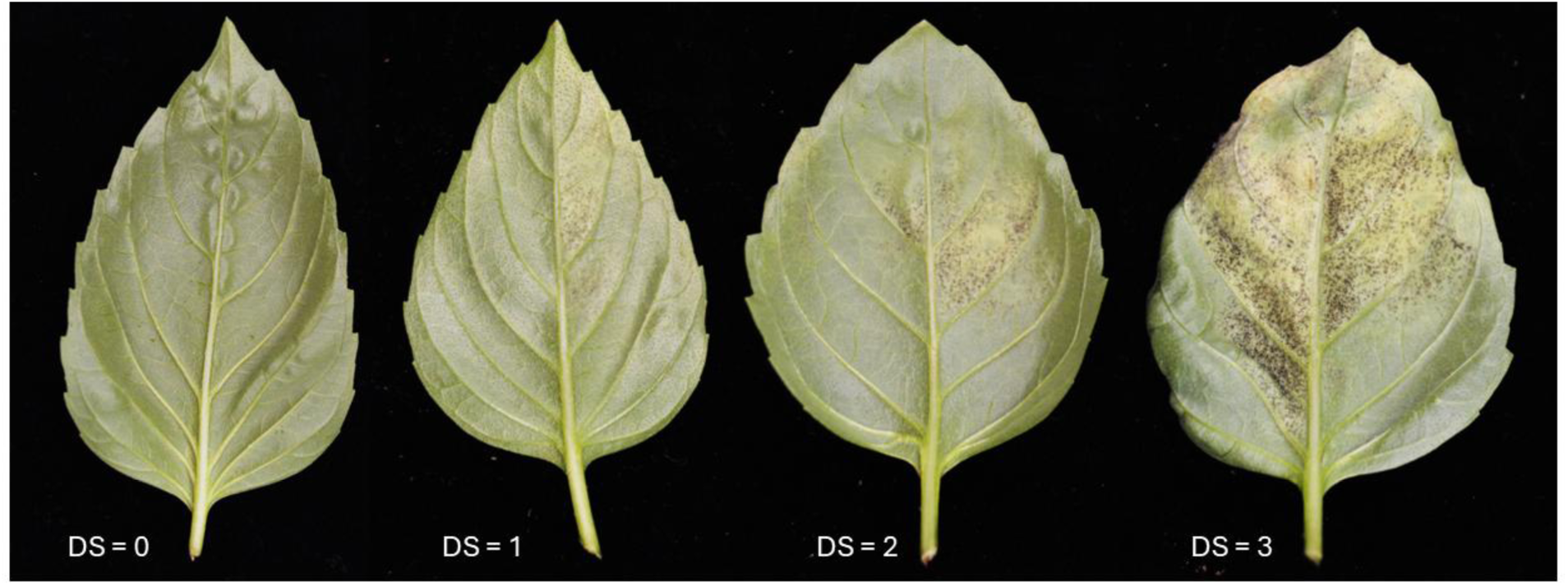
Four-point categorical scale used to rate the severity of Basil Downy Mildew (BDM) on the true leaves of inoculated plants. Examples of individual leaves showing DS scores 0=no sporulation, 1=low sporulation density (1-10% leaf area coverage), 2=moderate sporulation density (11-40%), 3=high sporulation density (41-100%).

### Data Analysis

Disease incidence data from each differential panel were analyzed using ANOVA to determine significant differences among cultivars, with means separated by Tukey’s HSD test at a significance level of ⍺ = 0.05. For disease severity data, which did not meet the assumptions of normality, the non-parametric Kruskal-Wallis test was applied, followed by Dunn’s post-hoc test to identify differences between groups. All analyses were performed in RStudio (Posit Team 2025).

## Results

### Establishment of *Peronospora belbahrii* inoculation and differential panel screening protocols

To objectively assess the interactions between diverse pathogen groups with diverse host genetic backgrounds, it is important to establish an environmentally controlled standardized protocol. Humidity incubation chamber, growing conditions, and *P. belbahrii* sporangia handling procedures were tested to find a highly reproducible protocol resulting in consistent infection of multiple whole basil plants in seedling trays or individual pots.

Humidity chambers were created with commercially available plastic bins containing 1-2 inches of standing water in the bottom (Supplementary Figure 1). Based on literature, the ideal infection conditions include a 24-hour post-inoculation incubation at 20°C in the dark, followed by a 20-25°C 12 h light/day cycle (Cohen et al. 2017). We found that the plastic bins were sufficient to maintain high humidity, typically ranging between 87% RH to 99% RH with an average of 96% in a greenhouse environment. These chambers provide adequate environmental conditions for *P. belbahrii* inoculation and incubation and can be disinfected easily after each round of experiments. All panel screens utilized a randomized complete block design to ensure that placement within the humidity chambers would have no impact on inoculation efficacy.

Sporulation requires a dew period with high humidity in the dark between 9 and 16 hours at 10 to 26°C (Cohen et al. 2017). We tested sporangia vitality by quantifying *in vitro* germination following collection, washing and storage procedures. Collection and washing (as described in Supplementary File 1) did not impact sporangia germination (Supplementary Table 1). Cryostorage of whole sporulating leaves can be utilized to maintain viable isolates, as described in other downy mildew pathosystems (Gill and Davidson 2005).

### Collection of *P. belbahrii* isolates

Three US isolates, five Israeli isolates, and one Italian isolate were tested in the differential panel screenings. An additional twenty-four unique *P. belbahrii* isolates were collected in the US from infected basil plants from 2018-2024. Of 27 total samples, 17 isolates were retrieved from commercially available downy mildew-resistant basil cultivars (Amazel, Prospera, Prospera Active, Devotion, Obsession, Passion, Thunderstruck), suggesting the rapid emergence of novel strains and/or race 1 in the US as early as 2018.

### Assignment of physiological races of *P. belbahrii*

Controlled-environment inoculations were performed to identify differential patterns of *P. belbahrii* virulence on a set of basil cultivars with unique sources of resistance. Panel screens of nine distinct isolates from the US, Israel, and Italy were tested, and performed in two locations when possible (Table 1).

**Table 1.**
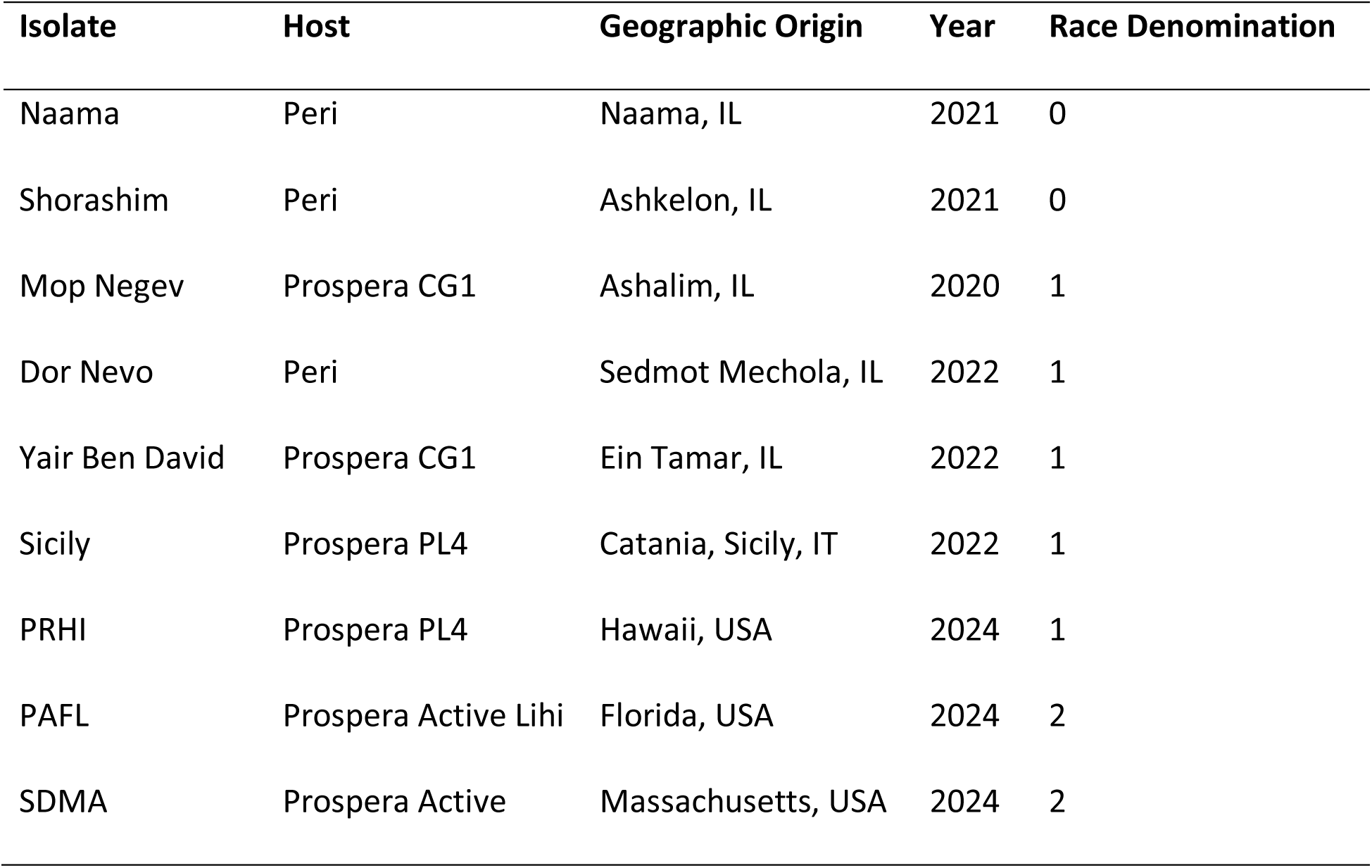
*Peronospora belbahrii* isolates screened at the University of Massachusetts Amherst and/or Rutgers University.

| Isolate | Host | Geographic Origin | Year | Race Denomination |
| --- | --- | --- | --- | --- |
| Naama | Peri | Naama, IL | 2021 | 0 |
| Shorashim | Peri | Ashkelon, IL | 2021 | 0 |
| Mop Negev | Prospera CG1 | Ashalim, IL | 2020 | 1 |
| Dor Nevo | Peri | Sedmot Mechola, IL | 2022 | 1 |
| Yair Ben David | Prospera CG1 | Ein Tamar, IL | 2022 | 1 |
| Sicily | Prospera PL4 | Catania, Sicily, IT | 2022 | 1 |
| PRHI | Prospera PL4 | Hawaii, USA | 2024 | 1 |
| PAFL | Prospera Active Lihi | Florida, USA | 2024 | 2 |
| SDMA | Prospera Active | Massachusetts, USA | 2024 | 2 |

Two isolates collected in Israel, referred to as Shorashim and Naama, exhibited virulence on susceptible Superbo and were avirulent against Prospera, Prospera Active series, *Ocimum* sp. accessions US86 and KL3 (Figure 2). The QDR lines Devotion, Obsession and Passion all demonstrated intermediate resistance or tolerance to Shorashim, with significantly reduced disease incidence and severity compared to susceptible Superbo. All Prospera and Prospera Active lines showed full resistance to the isolates, as did US86 and KL3. Naama exhibited similar virulence profiles across the panel, with slight differences in the QDR lines. Both Devotion and Passion were infected at significantly reduced levels (6.3-50% disease incidence) in replicated trials, but Obsession had a slightly higher infection rate. Replicated results confirmed that the overall responses across the differential panel were stable to define race 0 isolates, with the QDR lines conferring significant partial or full resistance, and the major DMR locus *Pb1* offering full resistance against both tested isolates. Prospera Active, US86 and KL3 lines exhibited full resistance to race 0 isolates.

**Figure 2.**
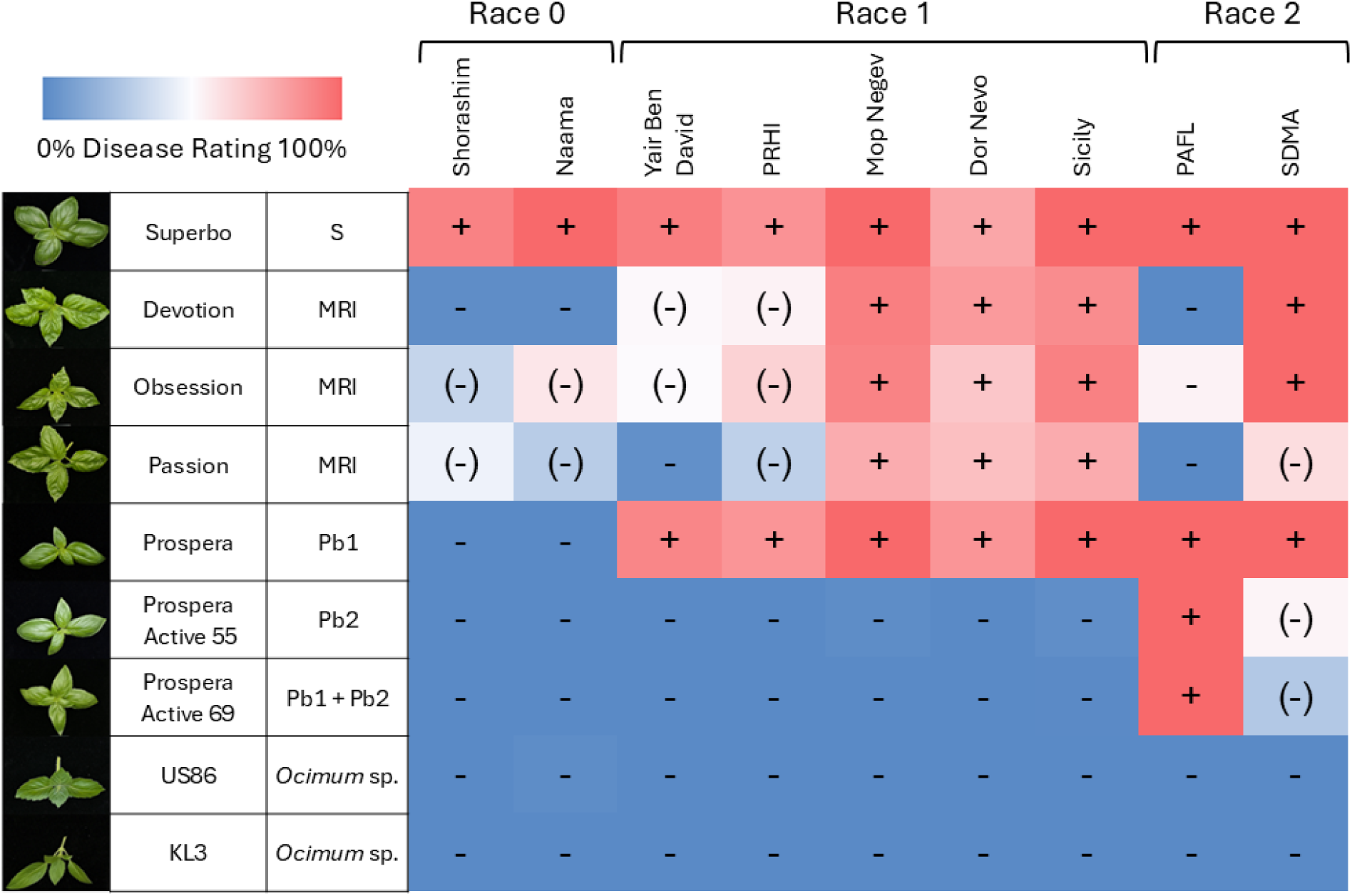
Differential Panel Summary. Heatmap of average disease ratings across differential panel lines tested against nine US and European *Peronospora belbahrii* isolates. Average normalized disease rating calculated: (Disease Incidence * Disease Severity)/Maximum Disease Severity. + denotes susceptible interactions, - denotes resistance, and (-) denotes intermediate resistance, defined as significant compared to both susceptible and *Ocimum* sp. lines.

The collection of race 1 isolates, first identified on Prospera material carrying the *Pb1* locus, includes Mop Negev, Dor Nevo, and Yair Ben David from Israel, Italian isolate Sicily, and HI from Pāhoa, Hawaii. The panel results confirm that the key shift from race 0 to race 1 is virulence overcoming the *Pb1* locus, resulting in severe infection on Prospera material comparable to that of highly susceptible Superbo. The Prospera Active lines, US86 and KL3 show full resistance to all race 1 isolates. The QDR lines displayed variable responses to race 1 isolates, showing intermediate resistance with significantly less disease than Superbo and Prospera in Yair Ben David and PrHI isolates, and susceptibility to isolates Mop Negev, Dor Nevo, and Sicily.

Isolate PAFL emerged as an outbreak on Prospera Active Lihi in a controlled environment production system in Florida in October of 2024. Differential panel screenings confirmed the unique virulence of this isolate on the Prospera Active lines carrying the *Pb2* resistance locus. This novel virulence pattern was used to characterize the newly identified Race 2 of BDM. Cultivars Superbo, Prospera and Prospera Active showed complete susceptibility to this isolate, with 100% disease incidence and heavy sporulation on all inoculated leaves (Figure 2).

Isolate SDMA was collected at the end of the outdoor growing season on Prospera Active Noga and Mia in a field trial in South Deerfield, Massachusetts, October 2024 (Scheufele et al. 2025). The infected leaves collected were nearing senescence after succumbing to heavy disease pressure in the field. The isolate was recovered and maintained on Prospera and Prospera Active plants before screening. High disease scores were collected from susceptible and Prospera lines, as well as QDR lines Devotion and Obsession. The Prospera Active lines showed moderate disease incidence and severity, and Passion showed comparable intermediate levels of infection.

In the differential panels used in this study, QDR varieties Rutgers Devotion, Obsession, and Passion exhibited partial resistance against at least one isolate of downy mildew. In many cases, this partial resistance was demonstrated with younger leaves exhibiting susceptibility, while older leaves remained asymptomatic (Supplementary Fig. 2). This finding challenges earlier research that suggested older leaves in susceptible basil varieties were more prone to higher rates of infection (Garibaldi et al. 2007). In these cases, the mechanism of quantitative disease resistance may be expressed as age-related, consistent with many QDR cultivars in other pathosystems (Coelho et al. 2009; DeMell et al. 2023; Hu and Yang 2019; Kus et al. 2002).

The emergence of race 1 and race 2 isolates are clearly delineated by *P. belbahrii* virulence on Prospera (*Pb1*) lines and Prospera Active (*Pb2*) lines, respectively. Continued isolate collection and screening may further separate race 1 into two groups based on the improved resistance of QDR accessions, as individual *P. belbahrii* isolates may display a unique virulence profile. The establishment of this panel provides a framework for screening environmentally active isolates, and further development of resistant cultivars and/or identification of new sources of resistant germplasm should be incorporated into the recommended differential panel accessions.

## Discussion

The practice of maintaining an active BDM isolate in a closed and controlled environment has advantages for continued experimental study. However, obligate pathogens are notoriously difficult to study due to their recalcitrance to axenic culturing. This research has resulted in the development, refinement, and utilization of replicable protocols for collecting and maintaining *P. belbahrii* isolates for large-scale studies. The developed inoculation protocol has been utilized for greenhouse cultivar trials, *P. belbahrii* isolate maintenance, and inoculation for DNA and RNA sample collections Other studies have described similar methods, in particular the maintenance of a *P. belbahrii* isolate on susceptible plants using a variety of high-humidity or dew chambers (Cohen and Ben-Naim 2016; Cohen and Rubin 2015; Pyne et al. 2014; Shao and Tian 2018). This differential panel and procedure has been refined and replicated in two locations with consistent results across the differential lines. Long-term storage and maintenance of viable inoculum also presents a key challenge for researchers to maintain multiple downy mildew isolates. We found that, consistent with other downy mildew pathosystems, sporulating leaves could be frozen and viable sporangia revived after several weeks to months. Downy mildew has been recovered from −80℃ freezing after years in some cases (Cohen et al. 2017; Withers et al. 2016). Our data suggest that partially dried sporulating leaves pressed against filter paper and frozen at a slower rate at −20°C improve viable sporangia recovery (Supplementary Table 1). The recommended best practices for long-term storage of *P. belbahrii* therefore, may include initial freezing of leaves to −20°C prior to deep freezing at −80°C.

These differential *P. belbahrii* isolate virulence and cultivar performance studies revealed the presence of multiple races of basil downy mildew in North America and the Mediterranean (for the first time). These trials complemented and confirmed results from various field trials, which have shown that, in some years and locations, different resistant cultivars have varied in their responses to BDM infection (Ben Naim et al. 2025; McGrath 2024).

The delineation of *P. belbahrii* race 0 and race 1 was first proposed in 2021, with race 0 described by avirulence on Prospera cultivars controlled by resistance gene *Pb1* and a unique SSR profile (Ben-Naim and Weitman 2022). A race 1 isolate was described by virulence against Prospera cultivars originally reported in New Jersey, with additional confirmation of a unique SSR profile (Ben-Naim and Weitman 2022), and our observations confirm the occurrence of this race in multiple years and locations in the US and Europe (Figure 2). The recent emergence of race 2 defined by virulence to the ‘Prosper’ *Pb2* resistance gene, in the Fall of 2024 in Florida and Massachusetts emphasizes the rapid selection that occurs in the evolution of the BDM pathogen following the introduction of novel sources of genetic resistance, and the continued need for improved BDM-resistant cultivars.

The intermediate resistance provided by the tested QDR lines (e.g., Rutgers lines) across races demonstrates the utility of incorporating multi-gene resistance into basil species where the BDM race population(s) are not known, or where it is known that multiple races of the pathogen are present. Across all isolates, at least one of the QDR cultivars conferred non-race-specific resistance, observed as a reduction in disease development and severity. In field settings, this ‘durable’ resistance is often overcome over periods of time and increasing pathogen pressure (Ben Naim et al. 2025). However, if early symptoms and signs of disease can be observed in both resistant and susceptible basils, additional interventions like chemical and cultural control practices can be quickly implemented to help reduce potential yield losses.

Ideally, combining QDR with major gene resistance in the future could be a strategy to improve durability in commercially viable sweet basil cultivars. Continued research into the genetic and environmental interactions governing these resistance mechanisms will be essential in shaping the future strategies for controlling basil downy mildew.

To our knowledge, this study is the first to formally describe two new basil downy mildew races using multiple *P. belbahrii* isolates from different geographic regions, establishing a differential panel to screen *P. belbahrii* isolates under controlled conditions. As new BDM-resistant sweet and non-sweet basil lines are developed, novel strains and new races of the pathogen will likely continue to develop. This study provides a framework based upon a new differential panel for race identification of existing isolates and classifying new *P. belbahrii* races as they emerge in the future.

## Acknowledgements

The authors would like to thank Meg McGrath, Susan Scheufele, Genevieve Higgins, DeWitt Thomson, Kai McClendon and Mordechai Schramm for assistance in procuring isolates. The authors also thank Chris Joyner and David O’Neil of the College of Natural Sciences Greenhouse at the University of Massachusetts for continued support with plant propagation.

## Supplementary Material

**Supplementary Table 1.** Evaluation of long-term frozen storage preparation on basil downy mildew sporangia viability.

| Storage Treatment | Temperature | % germination of sporangia recovered after 1 week |
| --- | --- | --- |
| Fresh sporangia (baseline germination) | - | 71% |
| Sporangia suspensions in 15% DMSO | -20°C | 2.7% |
|  | -80°C | 3.7% |
| Whole sporulating leaves in 50mL Falcon Tubes | -20°C | 29.7% |
|  | -80°C | 13.7% |
| Whole sporulating leaves pressed against sterile P5 filter paper in 50mL Falcon Tubes | -20°C | 49% |
|  | -80°C | 26.3% |

**Supplementary Figure 1.**
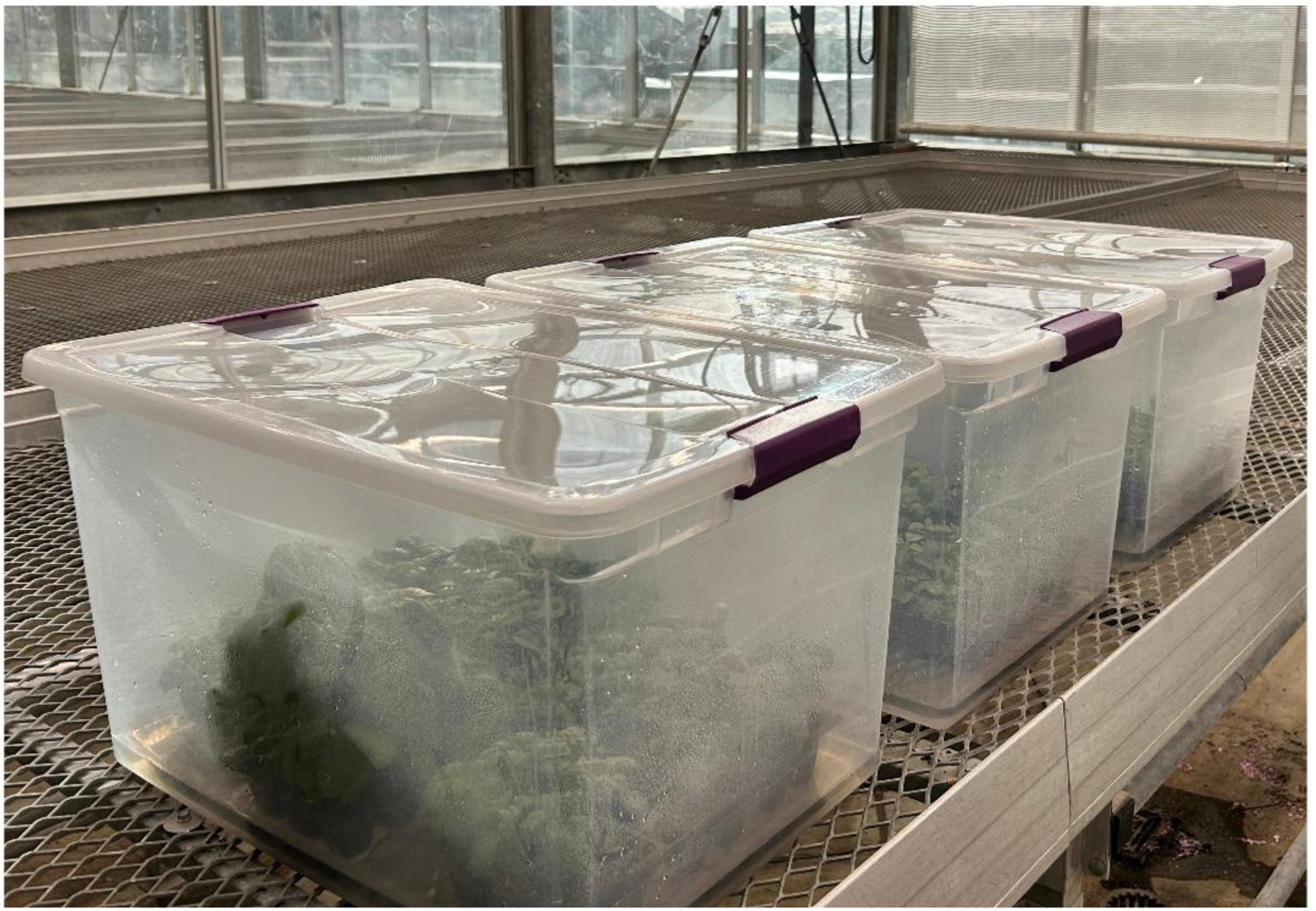
High humidity chambers used for basil downy mildew inoculations on a greenhouse bench (or growth chamber).

**Supplementary Figure 2.**
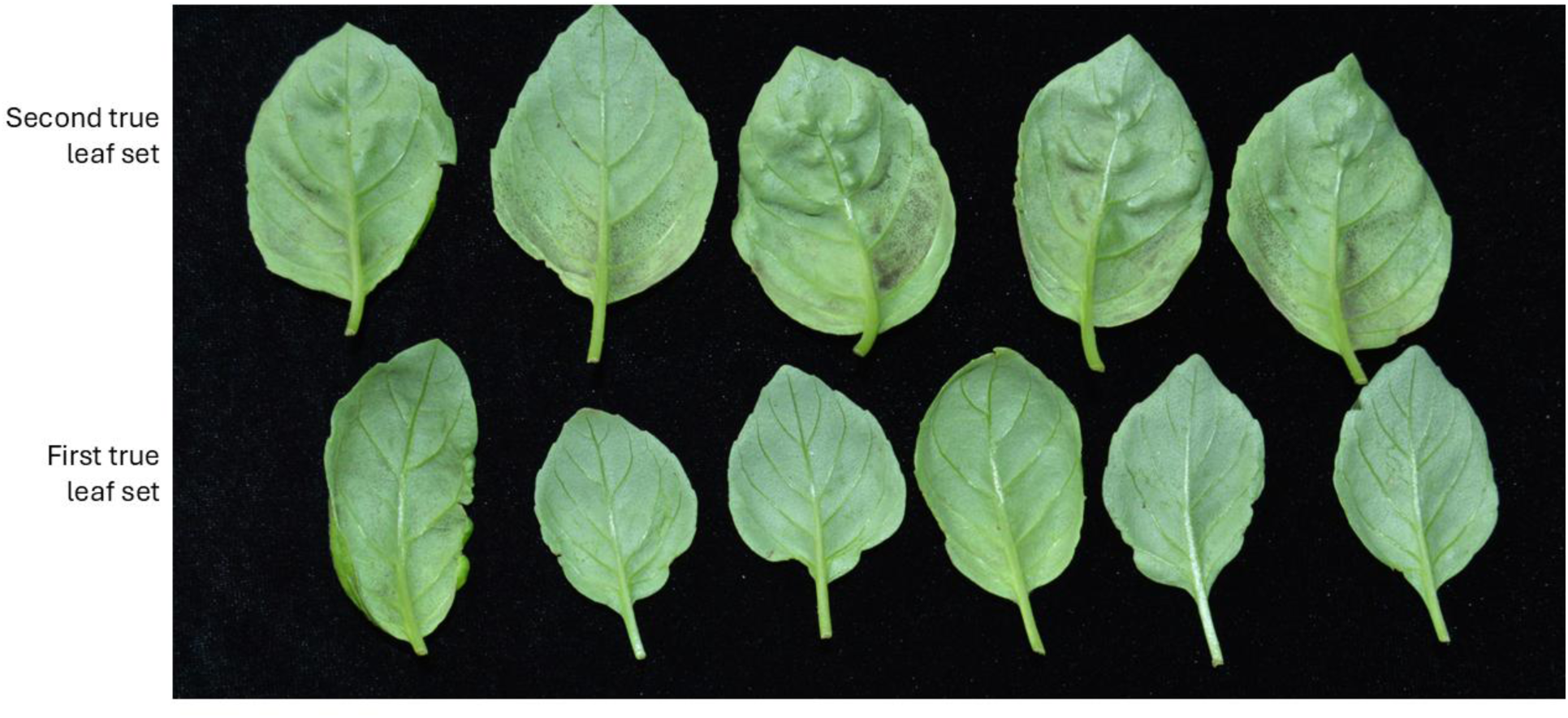
Quantitative resistant cultivar Passion leaves exhibiting partial resistance to Race 2 isolate SDMA. The second true leaf sets (top) showed more visible sporulation than the first true leaf sets (bottom), a phenotype that was observed across multiple isolates.

